# Antlions are sensitive to subnanometer amplitude vibrations carried by sand substrates

**DOI:** 10.1101/429209

**Authors:** Vanessa Martinez, Elise Nowbahari, David Sillam-Dussès, Vincent Lorent

## Abstract

European pit-building antlions (*Myrmeleon inconspicuus / Rambur 1842*) are studied in their capacity to detect vibrations generated by the locomotion of an ant (*Cataglyphis cursor*) outside the pit. These locomotions have been recorded by laser velocimetry and copied in detail in their time sequences. The sequences are replicated by micro-controllers digital outputs acting on piezoelectric transducers placed several centimeters outside the peripheries of the pits: their actions on the surface of a sand media create surface waves with particle accelerations that are 3 orders of magnitude less than *g*, alleviating any possibility of sand avalanche towards the bottom of the pit. Depending on the amplitude of the vibrations, the antlions answer back, generally by sand tossing. One remarkable feature is the time delay from the start of the cue and the aggressive behavior induced by this cue. This time delay is studied versus the cue amplitude. The result of this work is that anlions answer back within minutes to cues with amplitudes between 1 to 2 nanometers at the level of their mechanosensors, and within seconds to these same cues if they are preceded by a sequence of signals at the Ångström amplitude. This induced aggressive behaviour evidences the sensitivity to vibrations at extremely low level.

## 1. Introduction

The question of vibrational communication intermediated by mechanoreceptors in arthropods has received many attentions particularly in arachnids. Spiders have been the object of an extensive study (for a complete overview see the book of Friederich G. Barth (1)). The seminal work of Philip H. Brownell (2) in the seventies was devoted to the study of how desert scorpions locate their prey. The desert scorpions are poorly equipped with visual auditory and olfactory senses and hence their prey detection is by a large extend due to the vibration reception through their sensilla located at the tarsi. In terms of sensory systems, the antlion larvae is similar to the desert scorpion in its low ability in seeing and high sensitivity in vibration detection. In term of biotope, the antlion shares also similarities with the desert scorpion, at least for the perception of the vibrations: in both cases, those vibrations are propagated through a granular media. Unlike the desert scorpion, the hunting strategy of the sand digging antlion is to sit and wait for its prey. A funnel conical shape built by the antlion inside a granular media (generally sand) serves as a trap for any arthropod of reasonable size adventuring into it. Usually the presence of a prey creates an avalanche alerting the antlion hiding below a layer of sand at the bottom of the trap. The avalanche is maintained by the antlion tossing sand towards the prey by the intermediate of his eventually closed mandibles. The probability for a prey to escape from the pit is very low.

Since the senses of vision and chemoreception can be discarded (see D. Devetak in (3) chapter 16), we can associate the prey reception to sand movements either by avalanche or vibrations. The study explained hereafter focuses on vibration reception only. It is mostly inspired by the works of Le Faucheux (4), Fertin and Casas (5) and Mencinger-Vračko and Devetak (6) who identified the tufts of bristles around the body of the antlion as possible mechanoreceptive sensors. While the seminal work of Le Faucheux concentrate in identifying the bristles as mechanoreceptors by selectively ablating those, the independent studies in the groups of Casas and Devetak demonstrated the ability of the antlion to localize its prey in evidencing the strong correlation of sand tossing direction and prey direction.

Our study is made with no interference with the behavioural repertoire of the antlion, whereas the preys are replaced by artificial cues with a total control of the amplitude of the vibration: the vibration induced are far from originating any avalanche leaving only vibrations as a signalization of the presence of a prey. In this framework we specifically address the question of a threshold in the perception of vibration.

## 2. Material and Methods

### (a) Choice of animal model

The ecology of antlions informs us that 10 % of the species (7) are digging a pit in the sand or in a similar granular media. Once this construction done, the antlion waits at the bottom of the trap for a prey to descend the slope of this funnel. Depending of the size and weight of the prey, this last one might eventually be able to climb up the sand slope (8). On the other hand, the antlion, alerted by the sand avalanche and/or the sand vibration, tosses sand in the direction of the prey to aggravate the falling of the prey. We concentrate our study to the perception of vibrations by the antlion, alleviating any chance that this antlion is alerted by sand avalanche. *Euroleon nostras* has been studied extensively in the context of the mechanoreceptors physiology (4, 6, 9) and in the context of environmental factors and conditioning (10–12). Among the sit and wait antlions, *Euroleon nostras* and *Myrmeleon inconspicuus* are very similar in their behavior (13) therefore we considered that *Myrmeleon inconspicuus* antlions collected in the forest of Fontainebleau to be a good model for the studies concentrated to the mechano-perception of preys. The antlions selected for the study have been characterized in size and weight (see Supplementary Note-Subjects Table SN1).

### (b) Analysis of a natural cue inducing an agonistic behaviour of the antlion

It is usually evidenced that the antlion attacks the prey while it falls into its pit, and there is generally no such a behaviour when the prey is outside the pit. An agonistic behaviour is usually not demonstrated when the prey is outside the pit but some antlions, while they are in the process of digesting their prey nevertheless pay attention to their environment. It has been evidenced that an antlion is violently throwing away the caught prey if another ant is crawling at some distance (a distance up to 20 cm away from the border of the pit has been noticed in the laboratory). This last behaviour gives an hint about the mechano-sensory performance of an antlion. A first approach to quantify the amplitude of the signal received by the antlion would then be to measure the vibration signal propagating through the surface of the sand when this vibration is induced by an ant passing by. We made an attempt to measure these vibration in the following procedure: a *Cataglyphis cursor* ant, weighting 4 mg was placed around the pit of an antlion, while the antlion removed. At the bottom of the pit there is a 5 mm^2^ reflective stamp, adhesive on one side (that makes it embedded with the surface sand). The laser beam of the velocimeter strikes upon that small stamp. One films the ant in synchronization with the velocimetry recording. Given the noise level ^1^, no velocimetry signal can be retrieved while the ant is outside the pit. Inside the pit, two different types of signals are recorded, identified mainly by their amplitudes: while the ant crawls back to exit the pit some avalanches are created with amplitudes in the 1*µ*m − 10*µ*m scale and signals of amplitudes between 0.05 *µ*m to 0,5 *µ*m with frequencies around 1200 Hz, typical of the vibration propagation induced by a stepping function or an impulse on a sand surface. Since no quantitative measurement of a vibration amplitude triggering the attack of the antlion is possible by a direct monitoring of an ant walking around the antlion pit, it had been decided to mimic the induced sand vibration by an ant activity with monitored artificial hits on the ground.

### (c) Construction of an artificial cue

*Myrmeleon inconspicuus* is holomediterranean (14) and is very likely to share the same biotope with *Cataglyphis cursor*. Since *Cataglyphis cursor* develops a rescue behaviour (15) and is a possible model for co-evolution with an antlion predator (16) we choose to characterize its paces for further studies in these guidelines. Hence it is the exclusive source for the signal construction used in this study. The paces of a *Cataglyphis cursor* worker ant has been recorded by laser velocimetry (Fig.1). The ant crawls over a 10 cm × 5 cm paper surface illuminated from beneath by the laser beam of a commercial laser velocimeter (Polytech pdv-100). Velocimetry data has been acquired in conjunction with video screening in order to exclude data resulting from the eventual contact of the ant with the boundaries of the paper drumhead. Hereafter the selected velocimetry data has been replicated in the time intervals. A microprocessor board (Arduino Uno) is then coded to produce a 5 volt digital output pulse of 350 microseconds duration replicating these recorded time lapses. Whereas Fertin and Casas (5) choose to produce signals based on regular pulses of fragmented 4 kHz sinusoidal of 350 microsecond duration, we choose to apply 350 microsecond square pulses on the surface of the sand trough the action of the active surface of a piezo-electric transducer (PZT) in contact with the free surface of the sand substrate. We make use of ultrasonic PZTs resonant at 40 kHz (KPUS-40FS-18T-447) (see Materials and Methods). This PZT gives a flat response in the frequency range at which they are used (1/ (350 microseconds)), i.e. below the resonance frequency of the PZT. At the position where the antlion sits, the vibration is received with an amplitude ranging from 0.1 to 50 nanometers, depending on the output voltage and the relative position between the PZT transducer and the pit (Fig.2 and Supplementary Note-Signal calibration SN4).

**Fig. 1.**
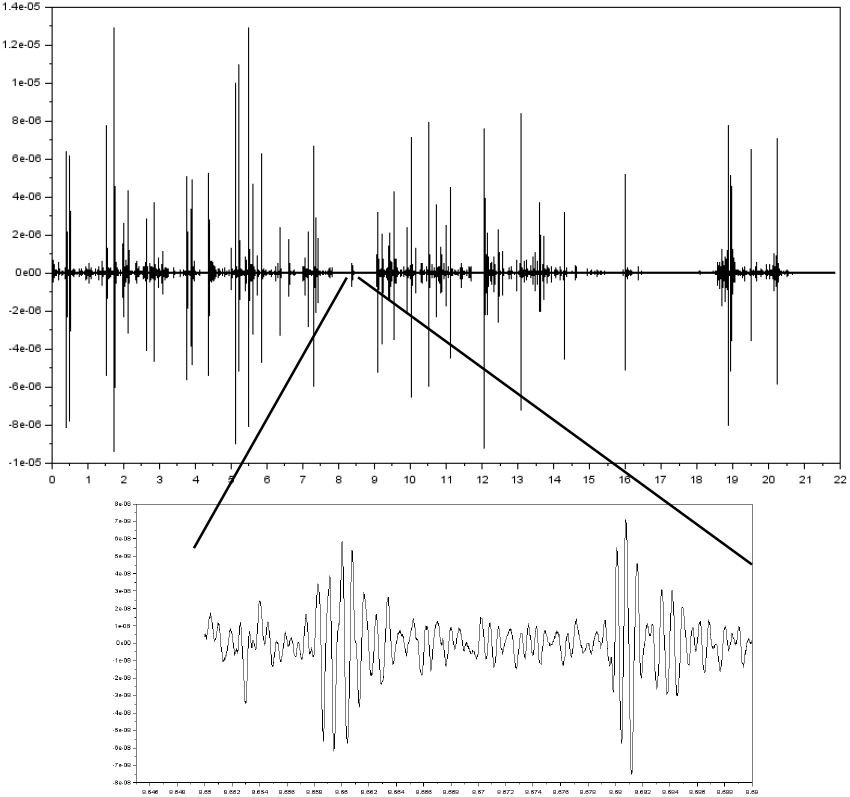
The figures above displays the transverse velocity of a 10 cm × 5 cm paper surface measured by laser velocimetry. This surface acts as a drumhead while a *Cataglyphis cursor* ant is walking on it. The vertical scale is in ms^−1^. The figure above is a velocimetry recording during 22 s. The figure below is a detail of this recording during 40 ms.

**Fig. 2.**
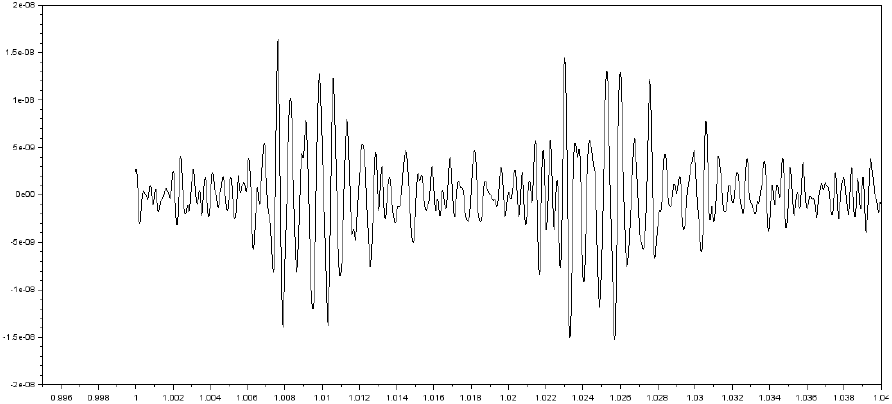
A PZT signal recorded a the bottom of an antlion pit. Two successive impulses of 350 *µ*s duration are separated by 20 ms time interval.

The device offers a reasonably small surface (10.4 mm diameter) in contact with the sand surface. Once the voltage applied on the PZT, the sand surface starts to oscillate with a frequency spectrum that depends on the mass of the object impacting the sand surface, the spring constant of the wire maintaining the PZT in contact with the sand surface and mainly on the elastic properties of the soil, as long the as the elastic regime is valid (17); the spring constant of the connecting wire is a parameter difficult to evaluate. The Fourier transform of the induced vibration by the impulse has been compared with the Fourier transform of the same impulse produced by a flat PZT inserted vertically in the sand and with the Fourier transform of the vibration produced by an ant walking on the sand surface. All frequency distributions are consistent (Fig.3).

**Fig. 3.**
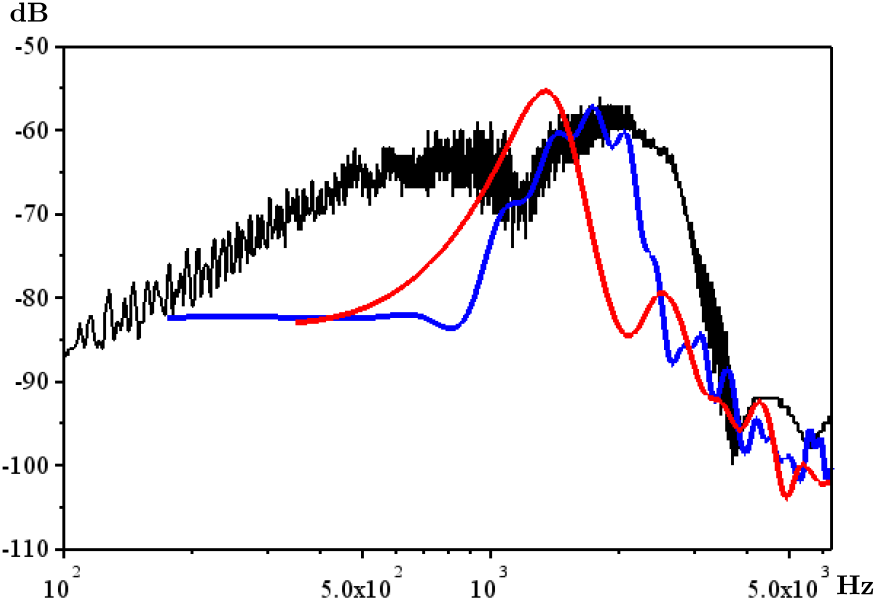
Fourier transforms of three vibration signals. The Y axis is the Fourier transform of the vibration velocity expressed in dB, while the X axis is the frequency in a logarithmic scale. Black: *Cataglyphis cursor* walking on the sand surface. Red: PZT with the vibrating surface leaning on the sand surface. Blue: flat PZT inserted vertically in the sand.

### (d) Implementation of the artificial cue and procedure

29 antlions (see Supplementary Note-Subjects Table SN1) have been tested with two independent set-ups. Each set-up consists of 8 independent containers individually filled with a 5cm thick layer of sand from the field where the antlions have been collected (Fig. 4 and Supplementary Note-Procedure). A PZT and an antlion are placed in each container. Each antlion has been fed exclusively with *Cataglyphis cursor* workers ant every 10 days. Tests have been performed outside feeding days. A coded Arduino Uno board powered by a dc battery, hence disconnected from a computer during the test, sends simultaneously the same digital output signal to the PZT’s in each of the individual containers. The signal is made of a pattern lasting 1.36 second copied from the laser velocimetry record of a *Cataglyphis cursor* walking on a paper surface (see Fig. 1 and Supplementary Note-Stimuli Fig. SN3) repeated continuously 250 times. These 250 period pattern is repeated two times with no time interval in between: one is at a given amplitude (phase 1), the other at another (phase 2). The microprocessor boards are coded to send the first impulse signal after a time interval chosen between 20 minutes to two hours. This preliminary silent time serves to discriminate the movements of the antlions inside their pits with those in correlation with the vibration cues. A camera (Sony HDR-XR200 VE - 4.0 megapixels or Sony Handycam HDR-CX740 - 24.1 megapixels) views this ensemble of 8 containers. In order to synchronize the time lapses between the camera and the micro-controller board, two photo-diodes shine in simultaneity, respectively with the signal in phase 1 and the signal in phase 2.

**Fig. 4.**
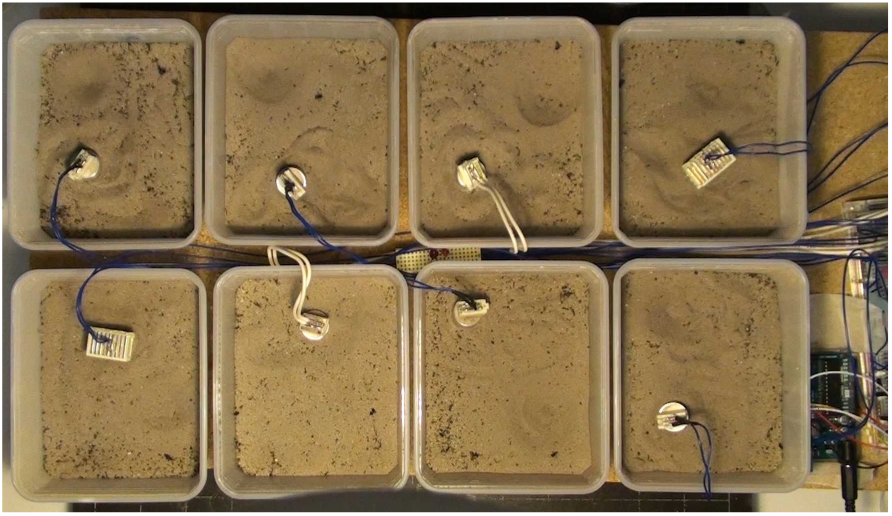
8 antlions in separated recipients are interrogated in parallel while submitted to identical cues.

These cue tests have been confronted to a control test consisting of presenting all the elements of the cue (actions of the microprocessor and the photo diodes without any voltage actually applied to the PZT’s). We noticed that 30 % of the antlions answer back to the actual signal while only 5 % respond to the control. A McNemar’s chi-squared test assigns a 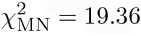 with a p-value = 1.083 10^−5^, thus rejecting the hypothesis of responses to idle signals.

## 3. Results

### (a) Spatial responses of the antlion to the artificial cue

The PZTs are placed in each container before the antlions are installed: hence the relative orientation between any particular antlion and the PZT is random. During the total duration of the experiment it has been noticed some re-localization of the antlions between each cue test. The antlions might also re-orient their body relatively to the PZT. It is worth to notice that, since the PZT offer a flat surface leaned on the surface of the sand, the vibration propagates with cylindrical symmetry. Hence one can characterize the complete geometry of PZTantlion relative positions by polar coordinates with the coordinate *r* being the distance between the antlion body and the PZT and the coordinate *θ* the angle between the antlion body orientation and the direction from the antlion to the PZT. Fig. 5 exhibits the occurrences of sand tossing in the angular *θ* repartition. Most of the time the antlions project sand in a direction lateral - and slightly backwards - to their body. This movement is consistent with the way the mandibles are used for projecting sand away in the action of pit building. Assuming that the sand tossing is towards the PZT these observed directions are also consistent with the observations made independently by Fertin & Casas (5) and by Mencinger-Vračko & Devetak (6).

**Fig. 5.**
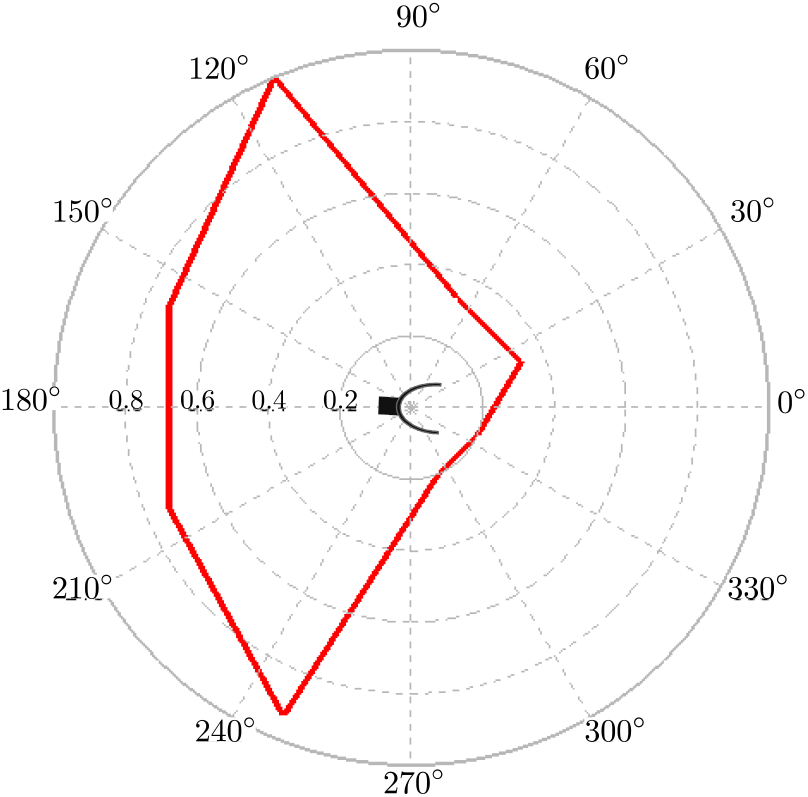
Repartition of the relative orientation of the antlion body with respect to the direction antlion to PZT emitter. The normalized distance from the centre represents the number of sand tossing in answering a cue test for a total of 67 tests. These sand tossing occurrences are collected in the angle distribution 0°−45°, 45°−90°, 90°−135°, 135°−180°, 180°−225°, 225°−270°, 270°−315° and 315°−360°.

### (b) Temporal responses of the antlion to the artificial cue

For what regards the sensitivity to vibration amplitude, this is addressed by the time delay between the start of the cue and the answer of the antlion to that cue. The following box-and-whisker diagrams (Fig.6) represents the delays of response of the antlions to the vibration patterns.

**Fig. 6.**
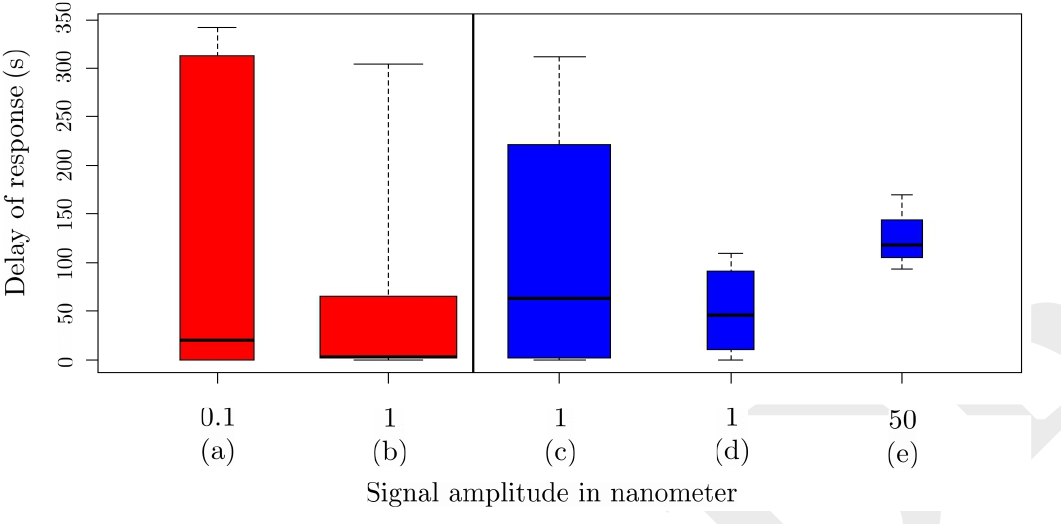
Box-and-whisker diagrams of antlion responses to an artificial cue as explained in the text. Each box shows the time response from the beginning of the cue. The bold horizontal line in each box is the median of the time delay. (a) cue in the range 0.1 nm lasting 340 s. (b) cue immediately following cue (a), in the range 1 nm lasting 340 s. (c) similar to (a) with an amplitude of 1 nm. (d) and (e) immediately follow (c) with a respective amplitude of 1 nm and 50 nm. The bases of each box is in proportion of the number of responses.

The boxes (a) and (b) are answers given by a group of 23 antlions. Box (a) is the time delay of response after the start of the signal, this signal being maintained during 340 s with amplitude varying between 0.06nm to 0.25nm depending on the recipient occupied by the antlion (see Fig. 4 and Supplementary Note-Amplitude of vibration Fig. SN5). Box (b) is the time delay of response counted from the start of the same signal with an amplitude 11 times higher, this signal appearing immediately after the signal (a) and lasting also 340 s.

The boxes (c), (d) and (e) are time delays for answers given by a group of 12 antlions to signals at higher amplitudes: from 0.5 to 3 nm in (c), to 0.8 to 5 nm in (d) and around 50 nm in (e).

The median of the time response of a 0.1nm amplitude signal (a) is 20 s after the start of the signal with 10 answers. The median of the time response of a 1 nm amplitude signal (c) is 63 s with 19 answers and the median of the time response of a 1 nm amplitude signal (b) with a 0.1 nm amplitude signal beforehand is 2.5 s with 34 answers. (c), (d) and (e) signals received 19, 4, 3 answers respectively. The medians of these last answers do not differ dramatically. Still, these data are essential to evidence by contrast that antlions who receive a preliminary signal of very low amplitude are more keen to react promptly a soon a signal of higher amplitude is present. The following histogram (Fig.7) is a test of validity of the assumption that the antlion is more prompt to react to a signal of moderate amplitude (1 nm) if this signal is preceded by a signal of very low amplitude (0.1 nm). The statistical model is a linear generalized mixed model with repeated measurement (GLMM).

**Fig. 7.**
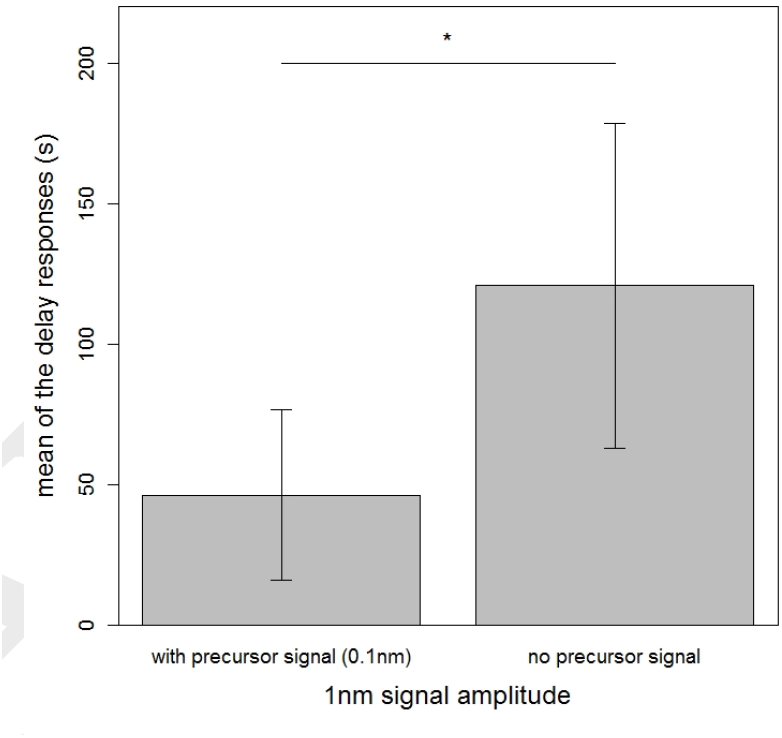
Bar chart presenting the mean value of time delay for the response of an antlion to a signal of 1 nm amplitude. left column is with a preceding signal of 0.1 nm amplitude (see text for explanation), right column is without a preceding signal. the p value of the GLMM test is 0,012.

## 4. Discussion and Conclusion

We sorted the antlion movements in three categories scaling up in aggressiveness intensity: partial exit of the hideout, mandible opening, mandible movements and sand tossing in the direction of the cue. At any level of behaviour we consider significant the first attack movement that follows the start of the artificial cue. We divided the artificial cue duration into several intensities and time sequences guessing that the antlion is making a choice with respect to the intensity of the cue. At high amplitude vibrations, from 1 to 5 nanometer amplitudes, the antlion takes minutes before showing a reaction. It leads to a simple interpretation: if the shape of the vibration may inform about the type of prey, its intensity should inform the antlion about its distance with respect of the center of its trap. A 1 to 5 nanometer amplitude signal and no sand avalanche means a prey outside or at the periphery of the trap and the time taken to react reflects on a possible balance between a chance of success of capture and the energy consumed in throwing sand towards this prey. The signal which consists into two sequences, the first 340 s with an Ångström amplitude followed by 340 s with a nanometer amplitude, is based on a guessing of an agonistic behavior. At first the prey seems far away from the trap and suddenly appears to be close. If the antlion is aware of the presence of a prey a some distance from its trap and attacks when it is worth doing it, i.e. when the prey approaches the trap, the behavior of a sudden response to the second part of the signal reflects this.

We emphasize that, in this experiment, the antlion behavior is the unique source of evidencing a vibration signal reception. Many arthropods have been studied for their ability to acquire information via substrate-borne vibrations, for a recent review see (3). Most of these physiological studies isolate the receptive organ to apply a mechanical stress and collect electrical signals. For instance, by these methods it has been demonstrated that the *Periplaneta americana* cockroach’s subgenual organ is a detector of vibration of exquisite sensitivity: mechanosensing experiments performed by means of elecromechanical cantilevers demonstrated a threshold of 0.2 nm (18) in the reception of the signal. These experiments will nevertheless not answer the question of how the animal perceives the signal. In that extend a behaviour physiology experiment such as the one presented here remains relevant in its neurophysiological aspects. The prolongation of our work will follow several directions: (i) basic physic and physiological considerations, that is, how the antlion reacts when submitted to signals propagating in different granular media with different phase and group velocities, (ii) characterization of pattern recognition by the antlion by the observation of its behavioural responses to different kind of time sequenced signals.

## Acknowledgements

We thank Pierre Tillier for indicating antlion locations in the forêt de Fontainebleau, Dušan Devetak for his precious help in the identification of the antlion species, Claudie Doums for providing *Cataglyphis cursor* ants, Paul Devienne for technical support and Karen Hollis for fruitful discussion. V.M. also acknowledges the support of the École doctorale Galilée - Université Paris 13.

## Supplementary Note

**Subjects**. 50 antlions have been collected end of May 2017 in the Fontainebleau forest in two different places; “95.2 la forêt des trois pignons” and “Notre Dame de Jouanne à la gorge du Larchant”. Among these, 29 antlions have been selected for the study. They are all instar 2 and 3 *Myrmeleon inconspicuus*. Their weights and sizes are summarized in the table SN1.

**Table SN1.**
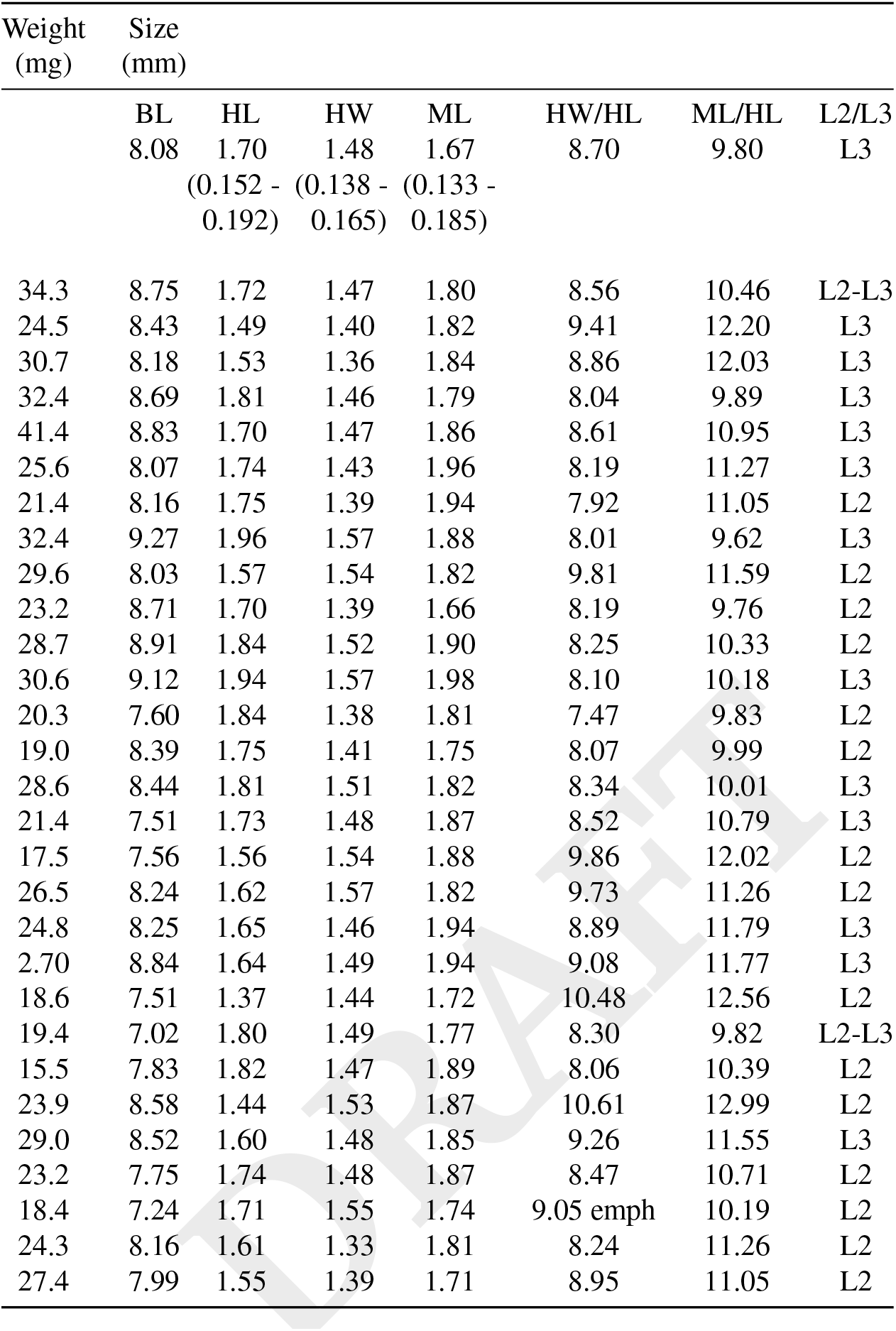
Weight and sizes of the *Myrmeleon inconspicuus* antlions selected for the study. BL: body length(excluding mandibles), HL: length of the head capsule, HW: head width, ML: length of the mandibles. The three first raws are data collected by Badano & Pantaleoni for the third instar L3 (19).

**Procedure**. The antlions have been placed individually in open containers filled with a sand bed layer 5 cm deep. Three sets of containers have been used: two sets with rectangular boxes as mentioned in the main text with dimensions 14 × 11 × 6.4 cm (L×W×H) and one set with circular boxes with dimensions 11.5 × 8.5 cm (D×H). All the experiments have been performed inside the laboratory without an anechoic environment. Hence it has been chosen to interrogate the antlions early in the morning or late in the evening. All antlions have been maintained in their collecting area sand. Fig. SN1 is an histogram of the granularity for the sand of the two locations of collection in the Fontainebleau forest. The particle size determination has been performed with a vibratory sieve shaker with sieves of mesh sizes 355,280,250,200,160,125,100 and 71 *µ*m.

Fig. SN2 is a picture of an antlion placed in a layer of sand in order to visualize the tufts of bristles (setiferous processes) attached on the lateral sides of the thorax (first, second and third segments on the picture). It clearly shows that the lengths and distances between the bristles are adapted for an optimum embeddedness of the sand particles between the seta.

**Fig. SN1.**
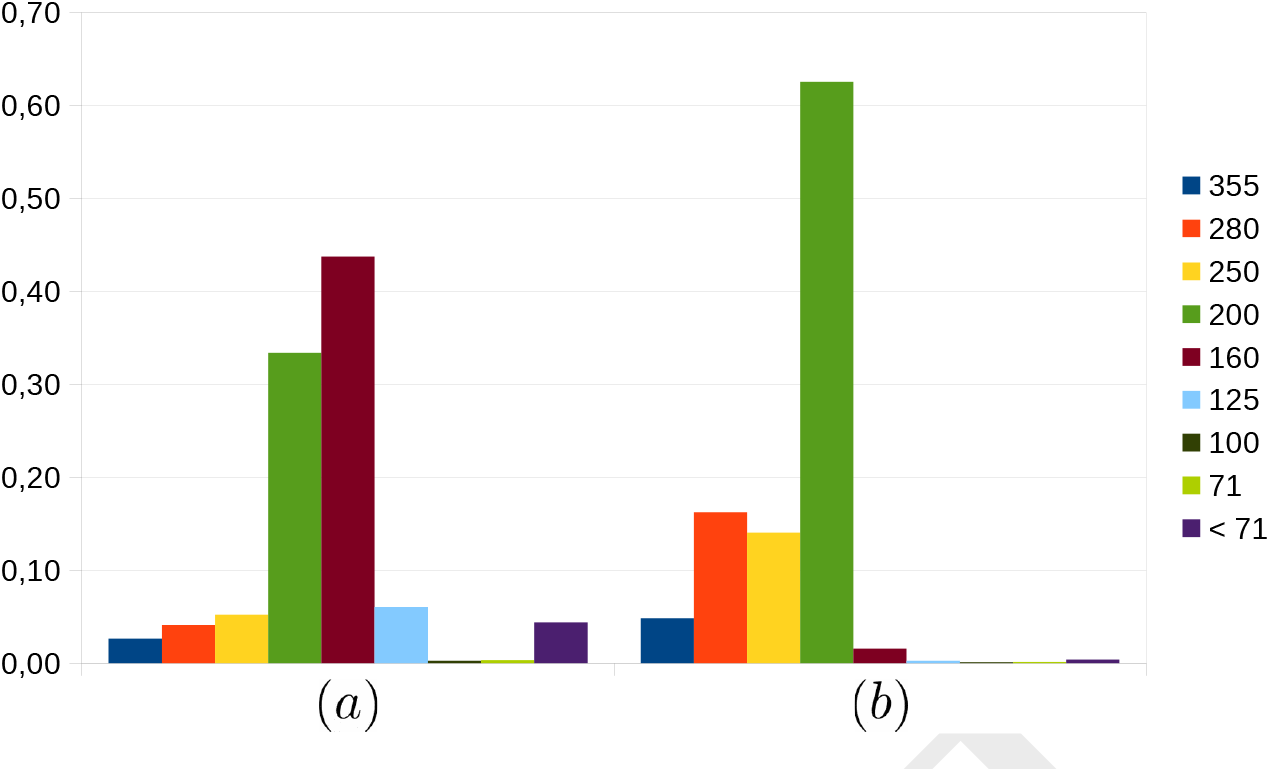
Determination of the grain size by a sieve shaker with calibrated sieve mesh sizes. Two samples of sand of weight ~ 1 kg each, collected at the respective named places in the forest of Fontainebleau (a) la cote 95.2 du massif de trois pignons, (b) La Dame Jouanne à la gorge du Larchant. The color code indicates the sieve mesh sizes in micrometers. Grain sizes are ordered from the biggest to the smallest.

**Fig. SN2.**
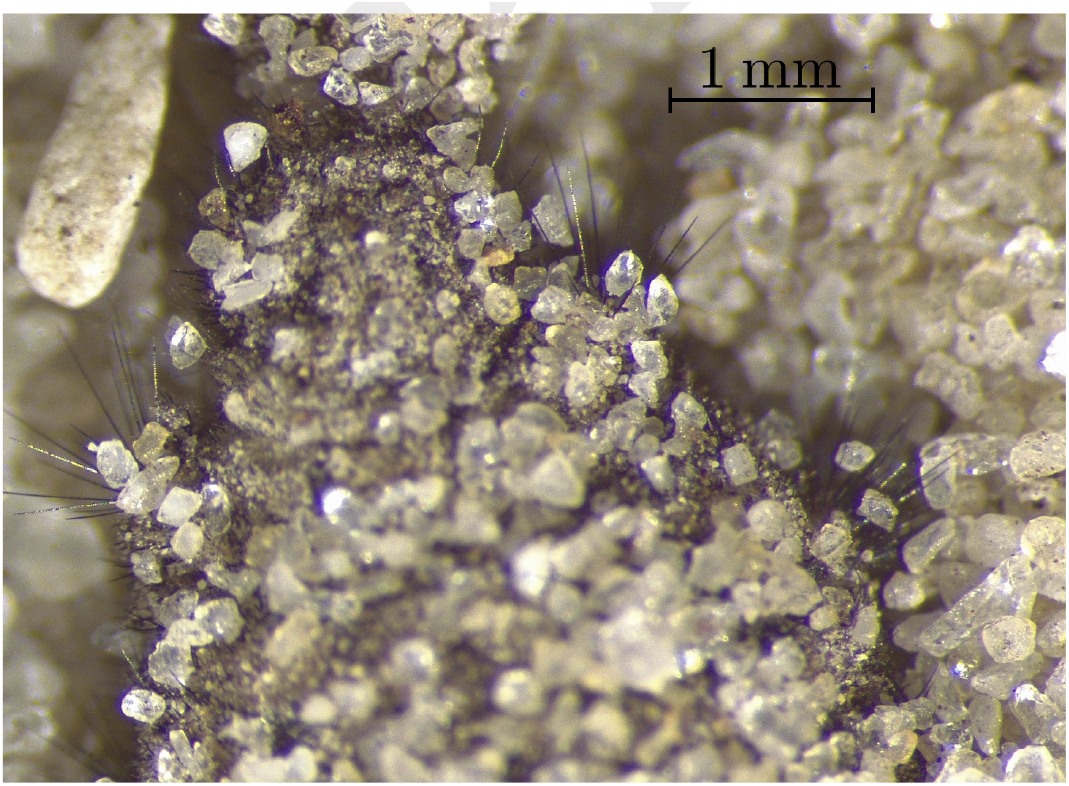
A portion of the thorax of a third stage larva of *Myrmeleon inconspicuus* laying at the sand surface. It displays the setiferous processes attached in the side of the first, second and third segments of the thorax.

**Stimuli**. An Arduino Uno microprocessor board is coded to produce an output voltage of 0 to 5 Volts with 350 *µ*s time on and a xx ms time off. The xx ms time off sequence is copied from a laser velocimetry record of a *Cataglyphis cursor* ant running on a paper bridge. On the average, the time off is 25.3 ms. An example of a time sequence is presented in Fig.SN3. It is produced from laser velocimetry recordings as described by the Fig. 1 where one extract a signal above a noise threshold yielding a bar code like signal.

### Signal calibration

- **Signal amplitude versus distance.** The amplitude of the signal is measured at the bottom of each sand pit fabricated by the antlions. In order to confirm that the measured amplitude by laser velocimetry is the proper one we verified that the vibration at the bottom of the antlion trap decreases with the distance from the PZT emitter. Fig. SN4 exhibits measurements of a vibration intensity with a ultrasonic sensor whose flat surface is leaning on the sand at several distances from the point illuminated by the velocimeter laser beam. These data are confronted to two different curve fittings. The blue line is a function ~ 1/*r*^2^ typical of a bulk wave; the red line is a function ~ 1/*r* typical of a surface wave propagation. The best fit is given by the red line indicating that one deals with surface waves. Since the source of vibration is given by a flat surface of diameter 14 mm while the wavelength of such surface wave is in the order 10 to 15 cm (20), one considers the vibration a such distances as far field.
- **Amplitude of vibration.** The laser velocimeter gives a calibrated measure of velocity. Knowing the sampling frequency of the velocimetry measurement (here 48 kHz) a numerical integration retrieves a vibration amplitude. In order to calibrate the output voltage of the microprocessor sending a signal to the PZT’s we measured the vibration at the bottom of each pit with a certain voltage. Considering the small signal amplitude, a periodic (350 *µ*s on − 5000 *µ*s off) signal is used for which the velocimetry signal is retrieved by an homodyne signal processing. Fig. SN5 is the repartition of vibration amplitude with a 2.5 V reference voltage of the digital output of the micro processor board.
- **Control of the voltage by a numerical potentiometer.** The amplitude is controlled by a numerical potentiometer. In the protocol of two consecutive cues, the first at the Ångström amplitude level, the second at the nanometer level (see main text), the output voltage of the microprocessor boards are 2.5 V and the potentiometer values are respectively set at 0 than 245: 0 value yields the maximum vibration amplitude whereas 245 value is less than 10 % of the maximum amplitude (see Fig. SN6).

**Fig. SN3.**
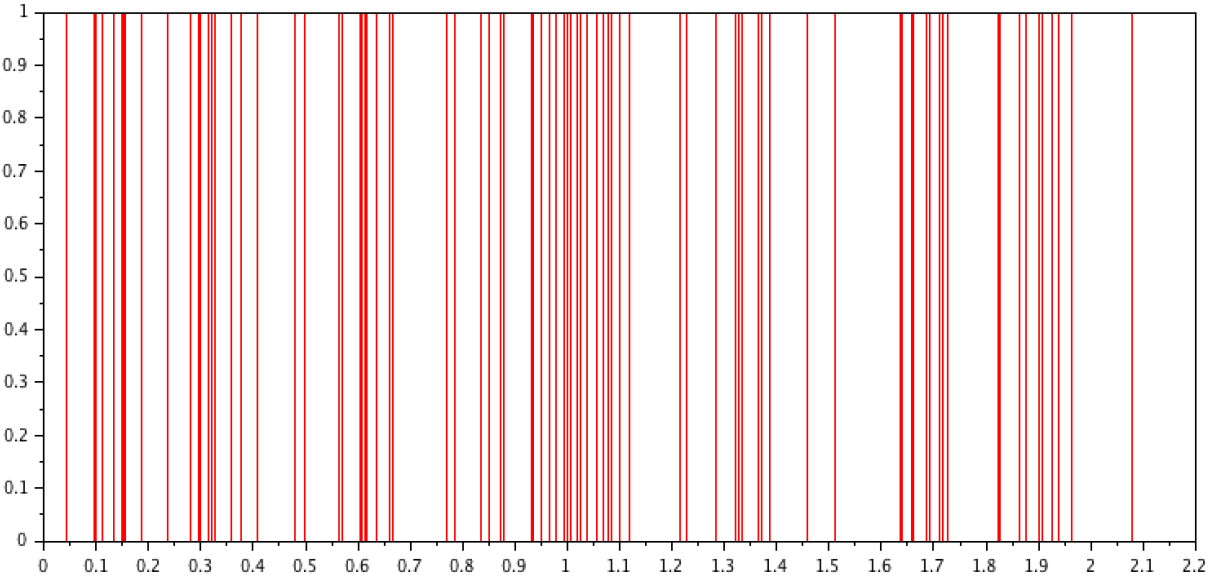
A 2.2 s sample of the laser velocimetry signature of a *Cataglyphis cursor* ant crawling on a paper surface as seen in Fig. 1. This signal is transformed into a bar code by sliding a temporal window over the square of the velocimetry data. By retaining as a signal the positive slope of this transformed data one obtains the starting moment of each impulse. This signal is at the source of the coding of the microprocessor.

**Fig. SN4.**
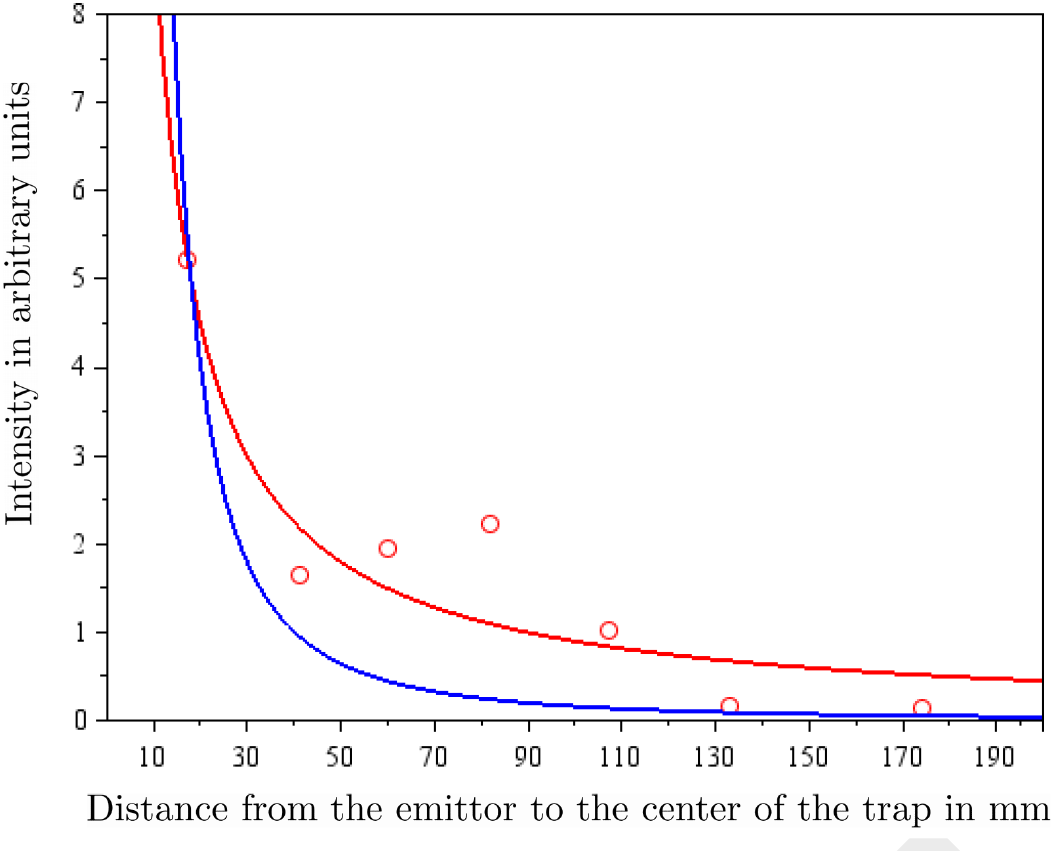
Intensity of the wave propagating through the sand as a function of the distance *r* between the emitter and the illuminated spot on the sand by the velocimeter laser beam. Red circles are data. Blue line is a curve fitting with a *a/r*^2^ function. Red line is a curve fitting with a *b/r* function. *a* and *b* are adjustable parameters. The red line is a better fit to the data suggesting that the wave propagates with an amplitude decreasing as 1/√*r*, hence with a decreased intensity ~ 1/*r*. The error coefficients for the curve fittings are: 2.19 for *b/r* and 7.6 for *a/r*^2^.

**Fig. SN5.**
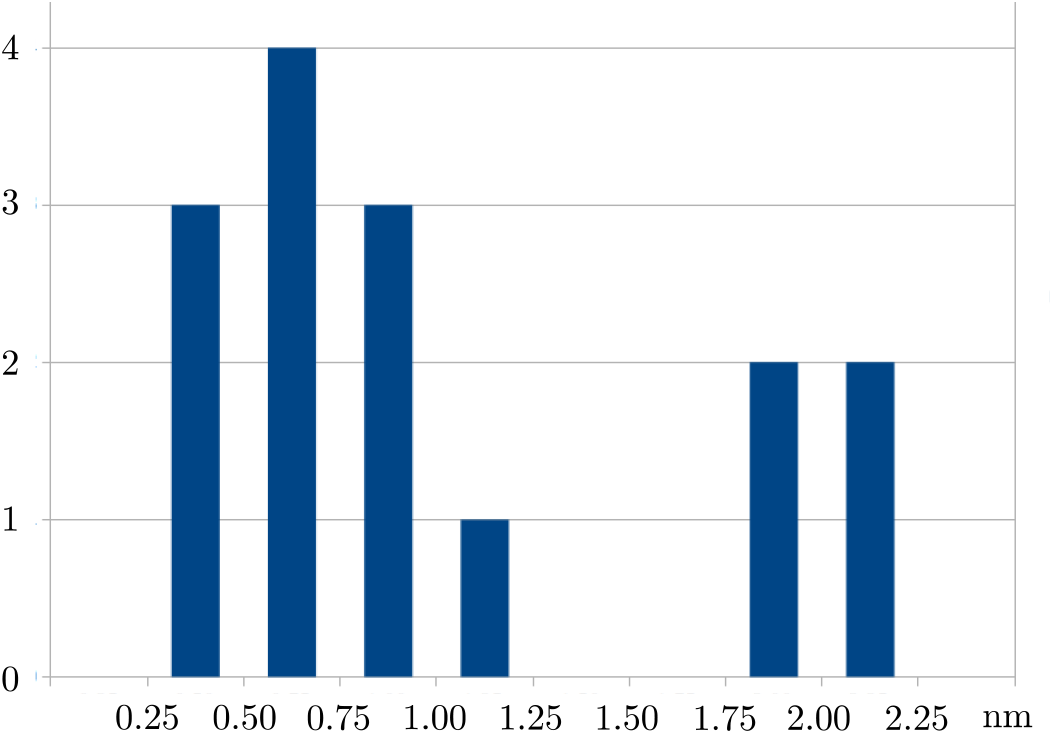
Repartition of the vibration amplitudes in each pit at the maximum output voltage of the microprocessor board. The vibrations below the nanometer level are essentially in the rectangular boxes where the distances between the antlion pit and the PZT are larger.

**Fig. SN6.**
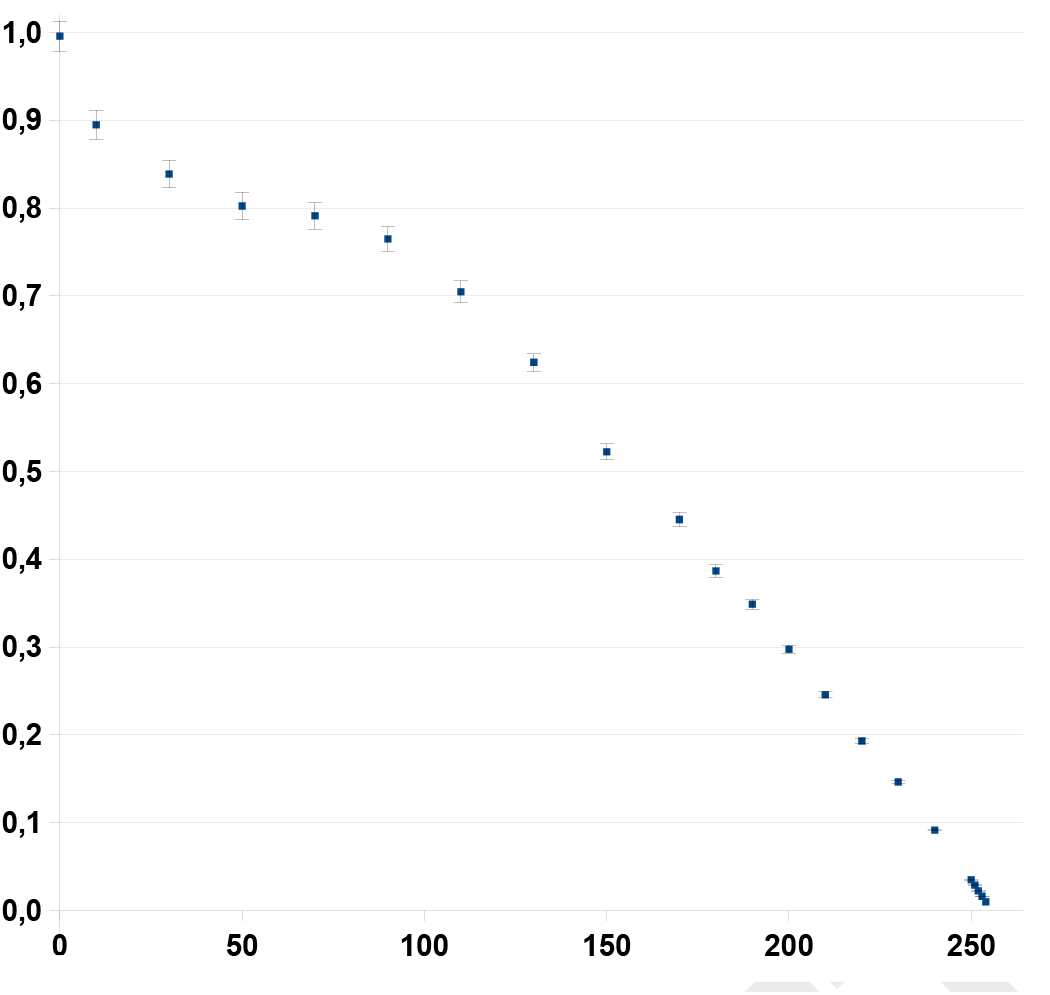
Amplitude of vibration measured by laser velocimetry at the tip surface of a PZT as a function of the integer value assigned to a digital 0−255 potentiometer.

**Chronology of the experiment**. In the experiment with one cue at 0.1 nm followed by 1 nm amplitudes 10 antlions have experienced 6 tests and 13 antlions have experienced 8 tests with the chronology of table SN2.

We have noticed a decay of responses to the cues with the repetition although the discrepancies in the chronology associated with each set (Fig.SN7).

**Table SN2.** Chronology of the two amplitude 0.1nm/1nm cue tests

| Test | 1 | 2 | 3 | 4 |
| --- | --- | --- | --- | --- |
| Set 1 | 17-10-18 | 17-10-19 | 17-10-23 | 17-11-07 |
| Set 2 | 17-11-07 | 17-11-08 | 17-11-09 | 17-12-01 |
| Set 3 | 18-03-14 | 18-03-19 | 18-03-21 | 18-03-23 |
| Test | 5 | 6 | 7 | 8 |
| Set1 | 17-11-08 | 18-03-13 |  |  |
| Set2 | 18-03-08 | 18-03-12 | 18-03-13 | 18-03-14 |
| Set3 | 18-03-08 | 18-03-28 | 18-03-30 | 18-04-03 |

**Fig. SN7.**
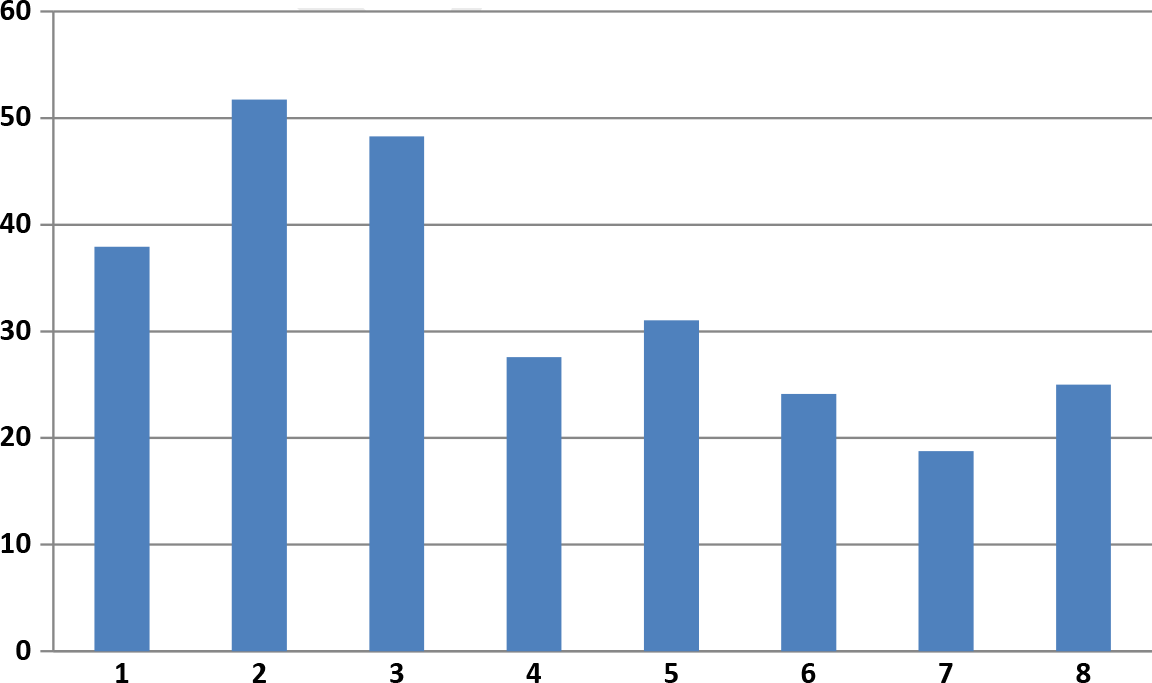
Percentage of antlion responses to the two amplitude 0.1nm/1nm cue tests.

## Footnotes

1 The velocimetry experiments are made with a commercial velocimeter Polytec PVD 100 with a velocity resolution 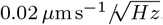. With a sampling frequency of 48 kHz the velocity resolution is ≃ 5 *µ*m s^−1^. We have been able to catch a signal as low as 0.8 nm at 1200 Hz, slightly above the noise limit.

## Bibliography

1. Friederich G. Barth. A Spider’s world, Senses and Behavior. Springer-Verlag, Berlin Heidelberg, 2002. ISBN 978-3-662-04899-3.

2. Philip H. Brownell. Compressional and surface waves in sand: used by desert scorpions to locate prey. Science, 197:479–482, 1977.

3. Reginald B. Cocroft, Matija Gogala, Peggy S.M. Hill, and Andreas Wessel, editors. Studying Vibrational Communication, volume 3 of Animal Signals and Communication. Springer Berlin Heidelberg, Berlin, Heidelberg, 2014. ISBN 978-3-662-43606-6 978-3-662-43607-3.

4. M. LeFaucheux. Le rôle des soies thoraciques dans la capture des proies par la larve d’euroleon nostras fourcroy (névroptère). Revue du comportement animal, pages 217–221, 1972.

5. A. Fertin and J. Casas. Orientation towards prey in antlions: efficient use of wave propagation in sand. Journal of Experimental Biology, 210(19):3337–3343, October 2007. ISSN 0022-0949, 1477-9145. doi: 10.1242/jeb.004473.

6. Bojana Mencinger-Vračko and Dušan Devetak. Orientation of the pit-building antlion larva Euroleon (Neuroptera, Myrmeleontidae) to the direction of substrate vibrations caused by prey. Zoology, 111(1):2–8, January 2008. ISSN 09442006. doi: 10.1016/j.zool.2007.05.002.

7. Johannes Gepp. Ameisenlöwen und ameisenjungfern. 2010.

8. Jérôme Crassous, Antoine Humeau, Samuel Boury, and Jérôme Casas. Pressure-Dependent Friction on Granular Slopes Close to Avalanche. Physical review letters, 119 (5):058003, 2017.

9. Dušan Devetak. Detection of substrate vibrations in the antlion larva, myrmeleon formicarius (neuroptera: Myrmeleonidae). Biol Vestn, 33(2):11–22, 1985.

10. Inon Scharf, Erez David Barkae, and Ofer Ovadia. Response of pit-building antlions to repeated unsuccessful encounters with prey. Animal Behaviour, 79(1):153–158, January 2010. ISSN 00033472. doi: 10.1016/j.anbehav.2009.10.017.

11. Lauren M Guillette, Karen L Hollis, and Audrey Markarian. Learning in a sedentary insect predator: Antlions (Neuroptera: Myrmeleontidae) anticipate a long wait. Behavioural Processes, 80:224–232, 2009. doi: 10.1016/j.beproc.2008.12.015.

12. Karen L Hollis, Heather Cogswell, Kenzie Snyder, Lauren M Guillette, and Elise Nowbahari. Specialized learning in antlions (Neuroptera: Myrmeleontidae), pit-digging predators, shortens vulnerable larval stage. PloS one, 6(3):e17958, jan 2011. ISSN 1932-6203. doi: 10.1371/journal.pone.0017958.

13. Maria Matilde Principi. Contributi allo studio dei neurotteri italiani. ii. myrmeleon inconspicuus ramb. ed euroleon nostras fourcroy. Bollettino dell’Istituto di Entomologia della R. Università degli Studi di Bologna, 14:131–192, 1943.

14. Pierre Tillier, Giacomino Matthieu, and Colombo Raphaël. Atlas de répartition des fourmilions de france. Revue de l’Association Roussillonaise d’Entomologie, Supplément au tome XXII, 2013.

15. Elise Nowbahari, Karen L. Hollis, and Jean-Luc Durand. Division of Labor Regulates Precision Rescue Behavior in Sand-Dwelling Cataglyphis cursor Ants: To Give Is to Receive. PLoS ONE, 7(11):e48516, November 2012. ISSN 1932-6203. doi: 10.1371/journal.pone.0048516.

16. Karen L. Hollis, Felicia A. Harrsch, and Elise Nowbahari. Ants vs. antlions: An insect model for studying the role of learned and hard-wired behavior in coevolution. Learning and Motivation, 50:68–82, May 2015. ISSN 00239690. doi: 10.1016/j.lmot.2014.11.003.

17. B. Andreotti, Yoël Forterre, and Olivier Pouliquen. Granular media: between fluid and solid. Cambridge University Press, Cambridge, 2013. ISBN 978-1-107-03479-2.

18. Stephen R Shaw. Re-evaluation of the absolute threshold and response mode of the most sensitive know “vibration” detector, the cockroach’s subgenual organ: A cochlea-like displacement threshold and a direct response to sound. Developmental Neurobiology, 25(9): 1167–1185, 1994.

19. Davide Badano and Roberto Antonio Pantaleoni. The larvae of European Myrmeleontidae (Neuroptera). Zootaxa, 3762(1):1–71, 2014. ISSN 11755334. doi: 10.11646/zootaxa.3796.2.4.

20. Philip H. Brownell and J. Leo van Hemmen. Vibration sensitivity and a computational theory for prey-localizing behavior in sand scorpions. American Zoologist, 41(5):1229–1240, 2001. ISSN 00031569.

